# Linear plasmids in *Klebsiella* and other Enterobacteriaceae

**DOI:** 10.1101/2021.12.14.472703

**Authors:** Jane Hawkey, Hugh Cottingham, Alex Tokolyi, Ryan R. Wick, Louise M. Judd, Louise Cerdeira, Doroti de Oliveira Garcia, Kelly L. Wyres, Kathryn E. Holt

## Abstract

Linear plasmids are extrachromosomal DNA that have been found in a small number of bacterial species. To date, the only linear plasmids described in the Enterobacteriaceae family belong to *Salmonella,* first found in *Salmonella* Typhi. Here, we describe a collection of 12 isolates of the *Klebsiella pneumoniae* species complex in which we identified linear plasmids. We used this collection to search public sequence databases and discovered an additional 74 linear plasmid sequences in a variety of Enterobacteriaceae species. Gene content analysis divided these plasmids into five distinct phylogroups, with very few genes shared across more than two phylogroups. The majority of linear plasmid-encoded genes are of unknown function, however each phylogroup carried its own unique toxin-antitoxin system and genes with homology to those encoding the ParAB plasmid stability system. Passage *in vitro* of the 12 linear plasmid-carrying *Klebsiella* isolates in our collection (which include representatives of all five phylogroups) indicated that these linear plasmids can be stably maintained, and our data suggest they can transmit between *K. pneumoniae* strains (including members of globally disseminated multidrug resistant clones) and also between diverse Enterobacteriaceae species. The linear plasmid sequences, and representative isolates harbouring them, are made available as a resource to facilitate future studies on the evolution and function of these novel plasmids.

**Significance as a BioResource to the community:** This study provides the first report of linear plasmids identified within the *Klebsiella pneumoniae* species complex and the first report in Enterobacteriaceae besides *Salmonella.*We present the first comparative analysis of linear plasmid sequences in Enterobacteriaceae, however whilst this family is highly clinically significant, the functional and/or evolutionary importance of these plasmids is not yet clear. To facilitate future studies to address these questions, we have publicly deposited (i) the collection of linear plasmid sequence data; (ii) isolates representative of each of the distinct linear plasmid phylogroups.

**Data Summary:** **The authors confirm all supporting data, code and protocols have been provided within the article or through supplementary information**

1. Whole genome sequence reads from *Klebsiella pneumoniae* isolates sequenced in this study have been deposited in NCBI SRA under the accession numbers listed in **Table S1**.
2. Representative annotated sequences of one linear plasmid per phylogroup have been deposited in FigShare, **doi 10.26180/16729126**.
3. A copy of all linear plasmid sequences that we assembled from publicly available genome sequence reads are available in FigShare, **doi 10.26180/16531365**. Read accessions for these are given in **Table S1**.
4. Eleven representative *K. pneumoniae* isolates harbouring linear plasmids described in this study have been deposited with the National Collection of Type Cultures (NCTC) and are available for purchase under the NCTC accession numbers listed in **Table S1.***K. pneumoniae* 1194/11 (representative of phylogroup B) has been deposited in the Microorganisms Collection Center, Adolfo Lutz Institute, São Paulo, Brazil. To request strain 1194/11 (IAL 3063, SISGEN ABBF09B), contact:

Microorganisms Collection Center
Culture Collection Laboratory
Instituto Adolfo Lutz, Sao Paulo State Department of Health
Address: Av Dr Arnaldo, 351, 10 floor, room 1020
Phone number: +55 11 3068-2884
Zip code 01246-000, São Paulo, Brazil
5. Alignments of terminal inverted repeat sequences for each phylogroup can be found in **Data S1**, available on FigShare, **doi 10.26180/16531371**.

## Introduction

Plasmids are extrachromosomal DNA that are frequently found in bacterial cells. The vast majority of plasmid molecules exist in a circular conformation, however linear plasmids have been found in several bacterial species, with the first description in *Streptomyces* in 1979 [1], and later in *Borrelia* [2], where they are universally present. A study of clinical *Enterococcus faecium* isolates recently reported the existence of a 143 kbp linear plasmid carrying a N-acetyl-galactosamine (GalNAc) utilization operon that could be transferred between strains via conjugation [3]. Linear plasmids appear to be exceedingly rare within Enterobacteriaceae, with the first, pBSSB1 (27 kbp), described in 2007 from *Salmonella* Typhi isolated in Indonesia [4]. Prior to this discovery, the only other linear replicons described within Enterobacteriaceae were those derived from bacteriophage, including pKO2 in *Klebsiella oxytoca* [5], N15 in *Escherichia coli* [6], and PY54 in *Yersinia enterocolitica* [7]. These bacteriophage-derived linear replicons are distinct from the true linear plasmids described in *Salmonella, Enterococcus, Streptomyces* and *Borrelia,* as they still possess bacteriophage-specific genes including those for the lysis pathways [5].

For replicons that are linear, there is a requirement to stabilise the terminal ends to ensure stability and appropriate replication, which in eukaryotes is achieved through the use of telomeres. In contrast, bacterial linear plasmids can either (i) create hairpin loops, as in *Borrelia* [8] and *Enterococcus* [3], or (ii) bind telomere-associated proteins to each end of the molecule with the assistance of terminal inverted repeats (TIRs), as in *Streptomyces* [9]. The *Salmonella* linear plasmid pBSSB1 was found to carry 1,230 bp TIRs with covalently bound proteins on the end, similar to *Streptomyces,* however these had no homology to any previously identified TIRs [4].

The *S.* Typhi linear plasmid pBSSB1 encodes two flagellar genes, an *fljA-like* gene and *fljB^z66^* [4]. *FljB^z66^* encodes the phase II z66 flagellin antigen, whilst the *fljA--like* gene is thought to encode the repressor of the chromosomally-encoded phase I flagellin antigen, allowing for phase II z66 antigen presentation [4]. Few other genes from the 27 kbp plasmid pBSSB1 have been characterised. Linear plasmids homologous to pBSSB1 have since been described in other *Salmonella* serovars, at a prevalence of ~0.3%, the majority of which carried the z66 flagellin genes [10].

In this study we report the discovery of multiple diverse linear plasmids in genomes belonging to the *Klebsiella pneumoniae* species complex (*K. pneumoniae* and six closely related taxa) within the Enterobacteriaceae. We demonstrate the linearity of these replicons using long-read and short-read sequencing, show they are reliably maintained within their natural host isolates during 10 rounds of laboratory passage, and identify homologs in the genomes of several other Enterobacteriaceae species. Clustering on the basis of gene content, we identify five major phylogroups of *K. pneumoniae* linear plasmids and describe their sequence characteristics in terms of size, GC content, TIR sequence and TIR length.

## Methods

### Identifying linear plasmids in K. pneumoniae species complex genomes

We screened for linear plasmids in the assembly graphs of 1,119 genomes of the *K. pneumoniae* species complex, including 452 from our own collection of human clinical and carriage isolates [11–13] and 667 publicly available read sets (see **Table 1**). Paired-end Illumina reads for each genome were assembled using Unicycler v0.4.7, using default parameters. The first assembly graph produced by Unicycler (001_best_spades_graph.gfa) was searched for the signature two-contig structure of a linear plasmid (a connected component of the graph consisting of one contig connected at both ends to the same end of another contig, see **Figure 1a**) using a custom Python script (available at doi 10.26180/16531374). We subsequently used these linear plasmid sequences as queries for a nucleotide BLAST search of the 1,119 genome assemblies, to recover instances where the linear plasmid sequence was present but had not fully assembled into the characteristic two-contig graph structure. This resulted in a total of 25 linear plasmid sequences, these have been deposited in FigShare, doi 10.26180/16531365.

**Figure 1:**
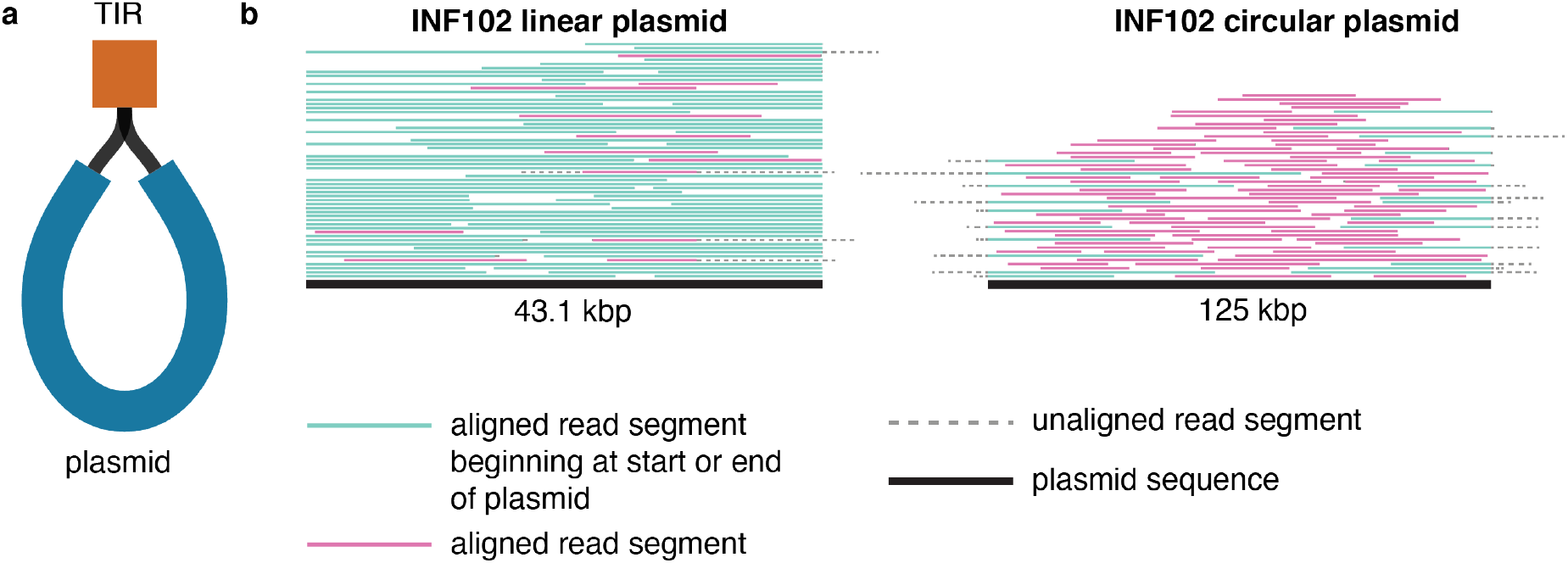
Using read sequence data to determine linearity of plasmid sequences. **a,** Short read assembly graph structure of a linear plasmid. The plasmid consists of two contigs, the main plasmid sequence (blue, labelled ‘plasmid’), connected to a second, shorter contig which is the terminal inverted repeat at both ends (orange, labelled ‘TIR’). **b,** Long reads aligned to the linear and circular plasmid sequences from INF102, with the total number of alignments shown capped at 100 to improve visualisation. The plasmid sequence is the thick black line at the bottom, and reads aligning to the plasmid are shown in green if the alignment starts at the beginning or end of the plasmid sequence, or pink if the alignment starts elsewhere. Segments coloured dotted grey indicate regions of the read that do not align. Alignments to the linear plasmid have very few reads which soft-clip off the ends of the plasmid sequence, indicating linearity. Conversely, alignments to a circular plasmid have many reads soft-clipping over the edges of the plasmid sequence, indicating that this replicon is circular.

**Table 1:**
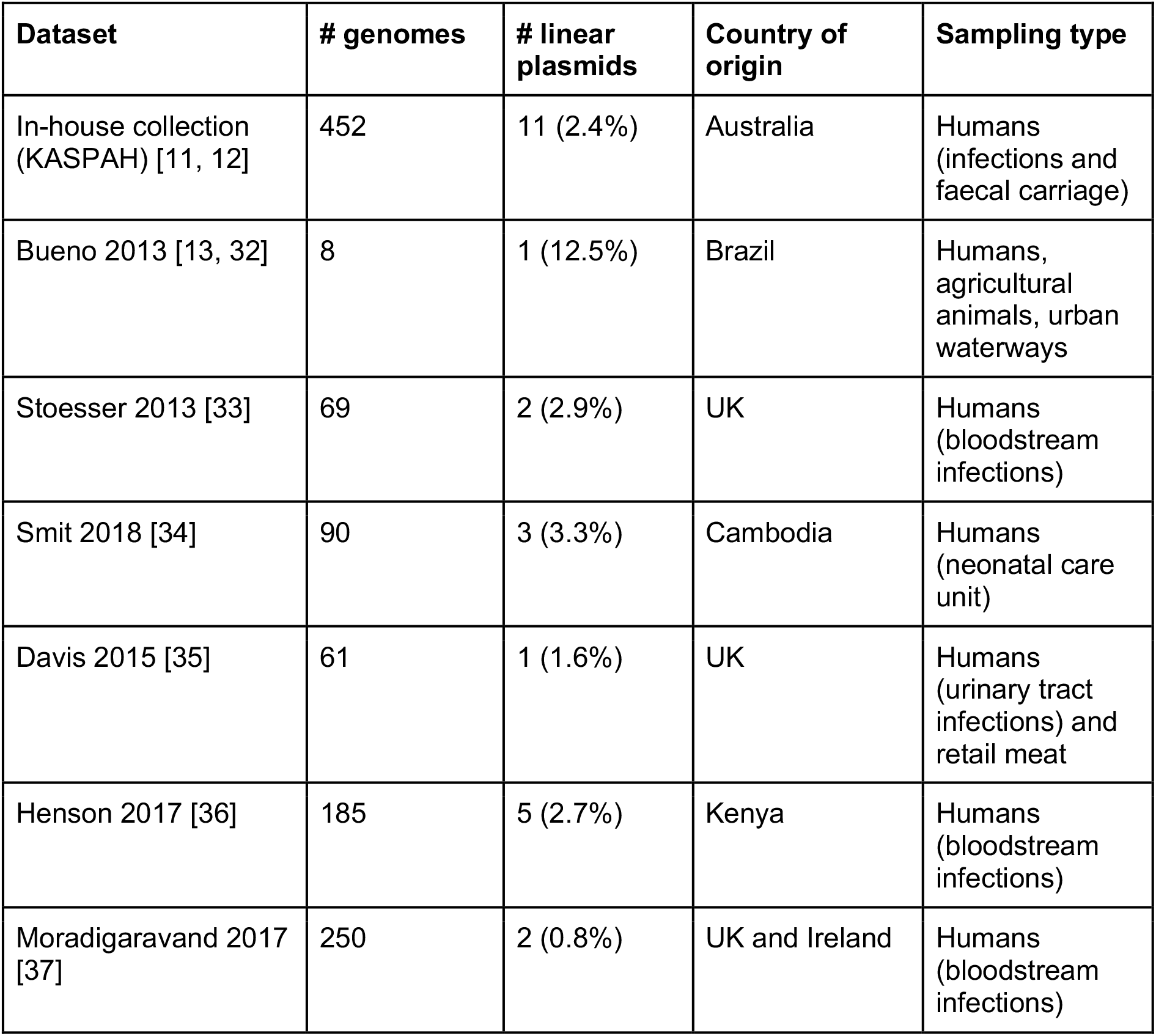
Number of genomes positive for a linear plasmid across multiple different studies from a variety of geographic regions and sampling types.

### Identifying homologs in other species

To detect homologous linear plasmids in other bacterial species, we performed a nucleotide BLAST search of NCBI (May 10th, 2021), using as queries each of the linear plasmid sequences identified in *Klebsiella,* as well as the pBSSB1 sequence (accession NC_011422). Hits with ≥90% identity and ≥60% coverage of a query sequence were considered as putative linear plasmid sequences (n=61). Metadata for each linear plasmid sequence and its host bacterium was pulled from the GenBank record for the corresponding whole genome sequence (WGS). To confirm the taxonomy and multi-locus sequence types (MLST) of the bacterial hosts of these putative linear plasmids, the chromosome sequence for each genome was uploaded to Pathogenwatch (https://pathogen.watch). For strain WP3-W18-ESBL-02 (in which plasmid 3, accession AP021975.1, was a hit to linear plasmid query pINF007 plasmid 3), Pathogenwatch was unable to detect a species, however the Genome-based Taxonomy Database (using GTDB-Tk [14] with database release 202, https://gtdb.ecogenomic.org) assigned it as a novel *Kluyvera* species, *Kluyvera ascorbata_B*. **Table S1** lists the species given by the submitter in GenBank, in addition to species detected by Pathogenwatch or GTDB, for all genomes.

### Plasmid stability analysis

For the 12 bacterial isolates in our collection with linear plasmids, we tested the stability of these plasmids during 10 passages in broth culture. Isolates from frozen glycerol stocks were streaked onto cation adjusted Mueller Hinton (CAMH) agar plates and incubated for 20 hours at 37°C. A single colony from each plate was streaked onto a fresh CAMH plate and inoculated into 3 mL of CAMH broth, and both were incubated for 20 hours at 37°C. From the broth culture, a glycerol stock and bacterial pellet, day 1 (D1) samples were prepared. This process was repeated 9 additional times to yield day 2-10 (D2-D10) samples.

Long-read sequencing (Oxford Nanopore Technologies, ONT) was performed as previously described [15]. Briefly, genomic DNA was prepared from the D1 and D10 bacterial pellets using GenFind v3 reagents (Beckman Coulter). A long-read sequencing library was prepared using the ligation library kit (LSK-109, ONT) with native barcoding expansion pack (EXP-NBD104 and NBD114, ONT). The library was run on a R9.4.1 MinION flow cell for 48 hours yielding 2.75 Gbp of reads. Reads were base called with Guppy v3.3.3 using the dna_r9.4.1_450bps_hac (high-accuracy) basecalling model.

To determine presence/absence and copy number of all plasmids in each genome, reads were mapped to their respective reference genome assemblies (listed in **Table S1**) using minimap2 v2.17 [16]. Mean read depth across each replicon in the assembly was calculated using the read alignments, and copy number for each plasmid was determined by dividing mean read depth across a plasmid replicon by the mean read depth across the chromosome.

### Confirming plasmid linearity

For the 12 linear-plasmid-positive isolates in our collection, reads from the day 1 (D1) ONT sequencing (see above) were aligned to their respective reference genomes using minimap2 v2.17 [16]. For each linear and circular plasmid sequence, we extracted all high quality read alignments (read identity ≥80%, alignment length ≥1,000 bp) that aligned within 90 bp of the end of the plasmid reference sequence. For these reads, we calculated the proportion that extended ≥100 bp beyond the edge of the plasmid reference sequence (and thus were soft-clipped ≥100 bp by the read aligner). If the replicon from which the reads originated was linear, we would expect to see few or no such soft-clipped reads, because the N’- and C’-terminal ends of the plasmid ssDNA molecules should match the start and end of the reference sequence (see **Figure 1b**). However, if the plasmid from which the reads originated was circular, we would expect to see many reads that are soft-clipped at the ends of the linearised reference sequence (see **Figure 1c**).

### Linear plasmid characteristics and relationships

To compare gene content across plasmid sequences, all 86 linear plasmid sequences retrieved from Enterobacteriaceae genomes were annotated using Prokka v13.3 [17], and genes were clustered into homologous groups using panaroo v1.2.4 [18], with a threshold of 70% amino acid identity to determine homology (details of clusters can be found in **Table S2**). The panaroo gene presence/absence matrix (**Table S3**) was subjected to hierarchical clustering using *hclust* in R (with default settings, i.e. Euclidean distance and *ward.D2* clustering algorithm) to generate a dendogram, which was cut into five phylogroups after visual inspection.

TIR length was calculated by taking each linear plasmid sequence, obtaining the reverse complement, and determining the length of sequence from the start of the forward and reverse complement sequences that were identical, with zero mismatches. Nearly all (except five) linear plasmid assemblies identified via nucleotide BLAST search of NCBI had very small TIRs using this method (n=56, between 0-54 bp). We assume this is the result of artefacts in the assembly process, which we are unable to explore without the underlying sequence reads; therefore plasmid sequences available only as publicly deposited assemblies without short reads were excluded from TIR length analyses. TIR sequences for the 25 linear plasmids generated from our assemblies were extracted, categorised into their respective phylogroups, and aligned using the clustalo algorithm in SeaView [19] to identify regions of homology within phylogroups (**Data S1**).

Nucleotide divergence between linear plasmid sequences was calculated by performing pairwise BLASTn alignments between all pairs of plasmids in the same phylogroup, and extracting the percent identity of the longest hit.

### Detailed annotation of representative linear plasmids

To further explore gene function in these linear plasmids, we undertook detailed annotation for one representative per phylogroup (A, INF019; B, 1194/11; C, INF102; D, INF007; E, INF352). Each representative was annotated using the RASTtk pipeline [20–22]. We screened for PFAM domains for all genes identified by RAST with hmmscan [23] via the EMBL-EBI server using default parameters. Resulting Pfam domains for genes with hits are listed in **Table S4**. To determine if any genes in the representative plasmids had homology to genes found in the Enterobacteriaceae, protein sequences were extracted from the RAST annotations and screened using BLASTp to the refseq_select database on NCBI, restricting results to Enterobacteriaceae. Genes with at least 50% protein identity to those in the Enterobacteriaceae were considered sufficiently similar to have a similar function. Representative plasmid annotations have been deposited in GenBank, accessions can be found in **Table S1**. To determine conservation of genes amongst plasmids in the same phylogroup, RAST annotations were matched with the Prokka annotations from the panaroo analysis.

### Trinucleotide profiles of linear plasmids and bacterial chromosomes

To investigate the potential donors of the linear plasmids into Enterobacteriaceae, we used *compseq* from the EMBOSS package [24] to calculate the frequencies of all possible trinucleotides in each of our 12 *Klebsiella* linear plasmids, their host chromosomes, as well as one representative per bacterial species (n=47,893) as defined by the GTDB database release 202 [25, 26]. We created a distance matrix using these frequencies with the *rdist* function in the R package *fields* (https://github.com/NCAR/Fields).

## Results and Discussion

### Identification of linear plasmids

We identified unusual structures in the assembly graphs of some *K. pneumoniae* in our in-house collection of genomes, which were consistent with linear plasmids with inverted repeats at either end (**Figure 1a, Methods**). We systematically screened for these structures in the assembly graphs of our in-house collection of *K. pneumoniae* species complex isolates, collected from human clinical infections or colonisation [11, 12] in an Australian hospital (n=452), as described in **Methods**. This screen yielded 11 genomes harbouring linear plasmids (2.4% of genomes) including seven *K. pneumoniae* and four *Klebsiella variicola* (**Table S1**). The corresponding isolates originated from nine patients, representing three instances of asymptomatic colonisation (*K. pneumoniae* ST359, *K. variicola* ST386 and ST642), one instance of simultaneous gut colonisation and pneumonia (*K. pneumoniae* ST37), and five instances of clinical infection (urinary tract infection with *K. pneumoniae* ST20, ST27, ST1449; wound infection with *K. pneumoniae* ST3073 and *K. variicola* ST347). The only extended-spectrum beta lactamase (ESBL)-positive (which confers resistance to the third generation cephalosporins) isolates amongst those with identified linear plasmids were two *K. variicola* ST347 isolated from the same patient nine days apart.

The linear plasmids were median 33,775 bp in size (range 31,739 - 44,271 bp), including the TIRs at either end. To confirm our hypothesis that these plasmids were indeed linear molecules, rather than the typical circular plasmid structure, we undertook additional sequencing using long reads, and aligned the long reads to each linear plasmid (see **Methods**). Plasmids were considered linear if there were few soft-clipped bases from reads aligned at the start or end of the linear reference sequence (unlike a circular replicon, where many reads are expected to overlap the ends of the linearised reference sequence, see **Figure 1b**). The 12 linear plasmids had a median of 3.5% (range 0.6-32.4%) soft-clipped start or end reads, compared to 98.5% (range 92.3-100%) for the circular plasmids (**Figure 1b, Fig S1, Table S1**). Additionally, all but two linear plasmids (those from *K. variicola* ST347) were supported by reads (median n=70, range n=10 to 177) that spanned the full length of the plasmid, including both TIRs (**Table S1**). Importantly, the soft-clipped parts of the reads did not map to the other end of the plasmid sequence (as would be expected for a circular plasmid), rather, they were chimeric reads, where two unrelated DNA segments have fused during library preparation [27].

To investigate whether other linear plasmids are present in the *K. pneumoniae* species complex, we generated and screened assembly graphs for an additional 667 publicly available read sets, which represent a diverse set of (mostly human clinical) isolates from multiple continents including Africa, Asia and Europe (**Table 1**). Across this set of genomes, we identified linear plasmid graph structures in an additional 14 genomes (2.1%, see **Table 1**). The corresponding isolates include 12 *K. pneumoniae* from humans (UK, Kenya, Cambodia, Brazil), one *K. pneumoniae* isolated from retail pork (USA), and one *Klebsiella africana* human blood isolate (Kenya).

Using as queries the sequences of the 25 linear plasmids that we identified from *Klebsiella* assembly graphs, we performed a BLAST search of the NCBI database to identify homologs in other genomes (see **Methods**). This revealed another 61 putative linear plasmid sequences; all were from Enterobacteriaceae, including *Klebsiella* (n=23, including 17 *K. pneumoniae), Salmonella enterica* (n=16, including pBSSB1), *Citrobacter* (n=8), *Enterobacter* (n=7), *Escherichia coli* (n=3), *Serratia marcescens* (n=2), *Phytobacter diazotrophicus* (n=1) and *Kluyvera ascorbata_B* (n=1) (**Table S1**). Genomes harbouring linear plasmids came from a wide variety of sources, including bacteria isolated from water (n=19), humans (n=13), food (n=4), animals (n=3), and plants (n=1) (**Table S1**). Amongst the linear-plasmid-positive *K. pneumoniae* were well-known carbapenemase-producing and ESBL producing clones: ST340 (n=3, KPC-4 and CTX-M-15), ST258 (KPC-2 and SHV-12), ST11 (n=1, KPC-2 and SHV-12), ST147 (n=1, OXA-181 and CTX-M-15). Hundreds of genomes of each of these clones are present in the NCBI database and the vast majority do not harbour linear plasmid sequences, suggesting that the linear-plasmid-positive variants are rare, and likely result from recent horizontal transfer but this has not resulted in clonal expansion during which the plasmid has been stably maintained. This is in contrast to the recent report in *E. faecium* where the linear plasmid *pELF_USZ* was stably maintained in a host lineage during >2 years of clonal spread in a hospital [3].

### Characteristics of linear plasmids in Enterobacteriaceae

We compiled the full set of 86 linear plasmid sequences (25 identified from assembly graphs, plus 61 inferred from homology via BLAST) and clustered them by their gene content (see Methods). This revealed five distinct linear plasmid phylogroups (which we labelled A-E, see **Figure 2, Table S2 & S3**), with very little gene sharing between phylogroups (genes defined as homologous if they had >70% nucleotide identity). Each phylogroup included sequences from multiple genera, notably all five phylogroups were detected in both *Klebsiella* and *Salmonella* (**Figure 2**). No genes were present across more than two phylogroups, but each phylogroup had a core set of genes found in ≥95% of plasmids in that group; these represented between 15% (phylogroup E) to 47% (phylogroup B) of all genes found in that phylogroup (**Figure 3a, Figure 4a**). Nucleotide diversity within phylogroups varied (**Figure 3b**), with phylogroup A displaying significantly greater pairwise divergence across the full plasmid sequence than phylogroups B, C and D (mean 6% divergence vs mean 2.6-4.2%, p<1×10-16 using Wilcoxon test for A vs B, C or D). Phylogroup E showed a high range in divergence (0-16%, mean 4.2%), due to the presence of two divergent subgroups (see **Figure 2**).

**Figure 2:**
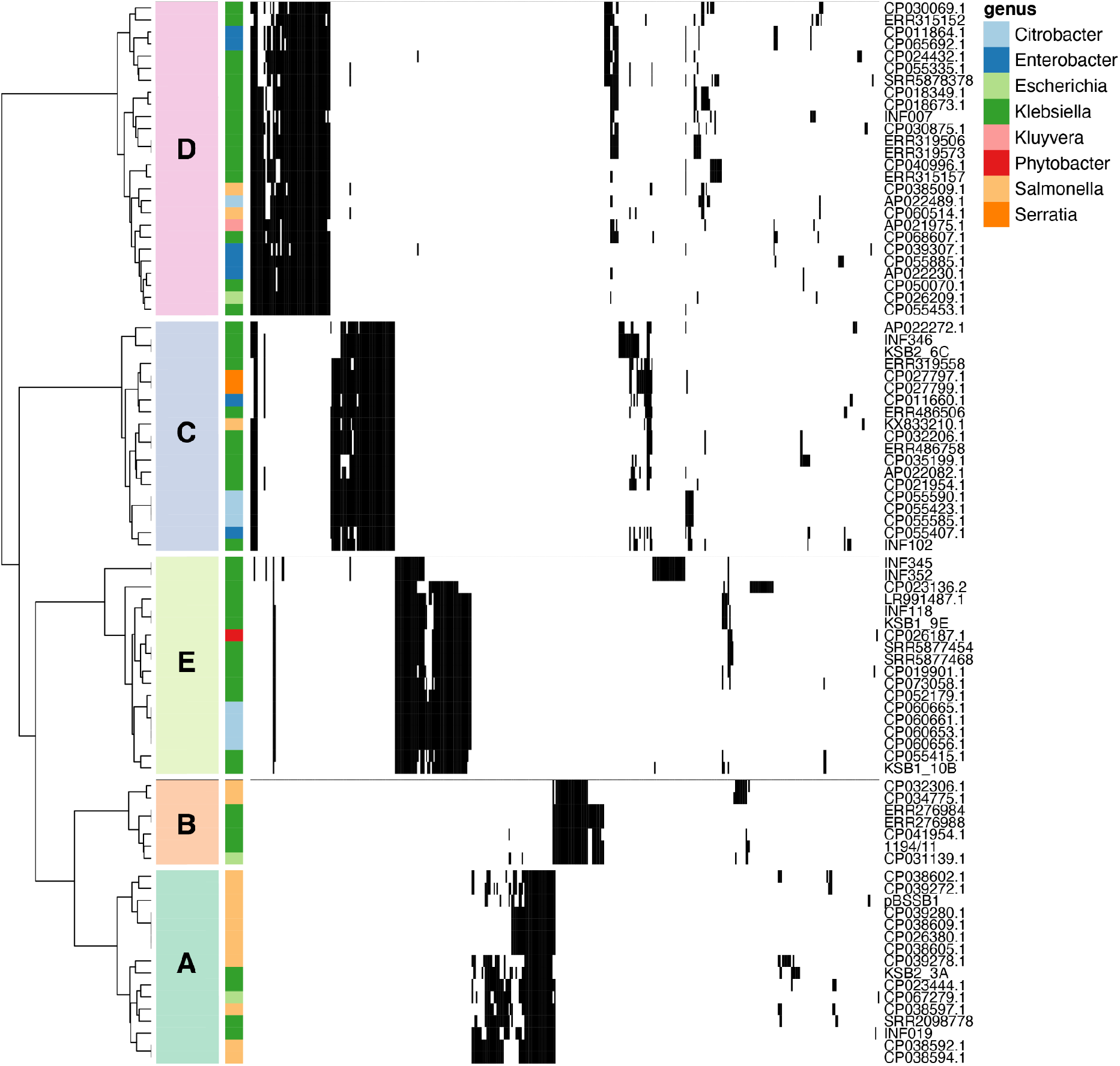
Hierarchical clustering of linear plasmids based on gene content. Plasmids were clustered with the *hclust* algorithm using the *ward.D2* method, and divided into five phylogroups (labelled in coloured boxes). Rows are annotated with the bacterial genus each linear plasmid was found in as per legend. Black indicates the presence of a gene, white absence. Plasmids are labelled with their names as per **Table S1**, and details of each gene can be found in **Table S2**.

**Figure 3:**
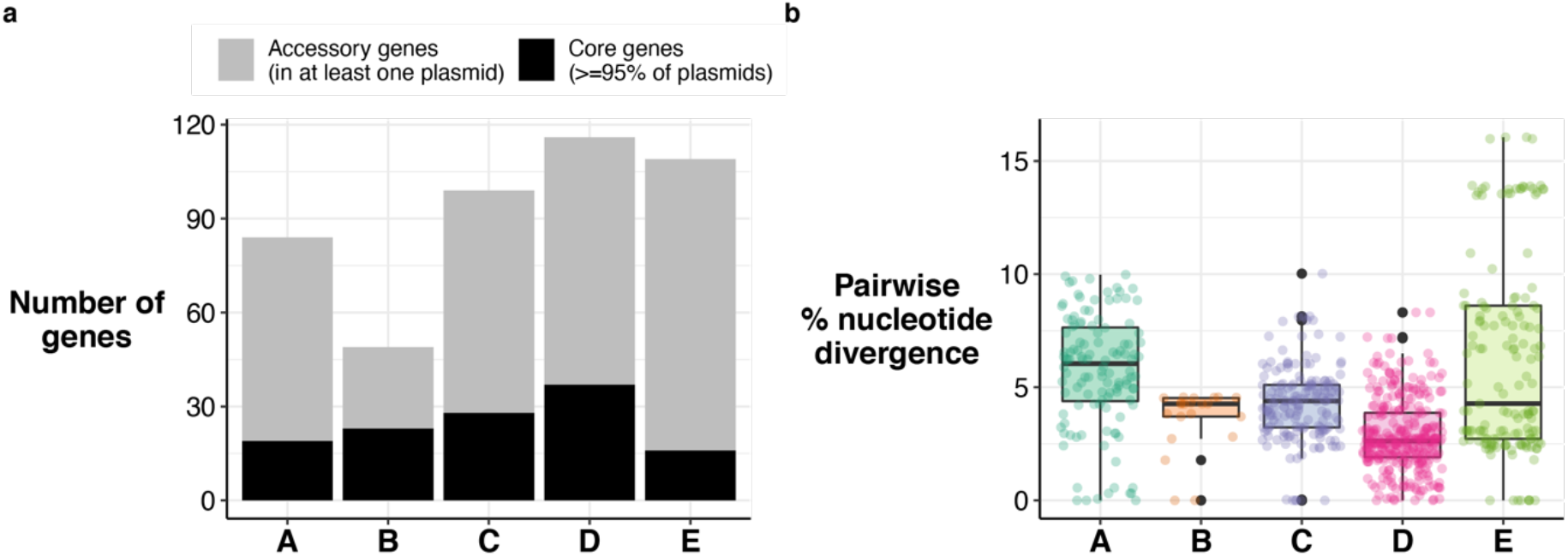
Core and accessory gene content by phylogroup, and nucleotide divergence by phylogroup. **a,** Number of core and accessory genes in each phylogroup. Bar height indicates the total number of genes found in at least one linear plasmid in each phylogroup. Black indicates the number of core genes (found in ≥95% of plasmids); grey the number of accessory genes, as per legend. **b,** Distribution of pairwise nucleotide divergence within each phylogroup. Boxplots show median (thick black line), 1^st^ and 3^rd^ quartiles (edges of box), and solid lines give 1.5x the interquartile range. Outliers are shown as black dots. Individual values are shown as coloured dots.

**Figure 4:**
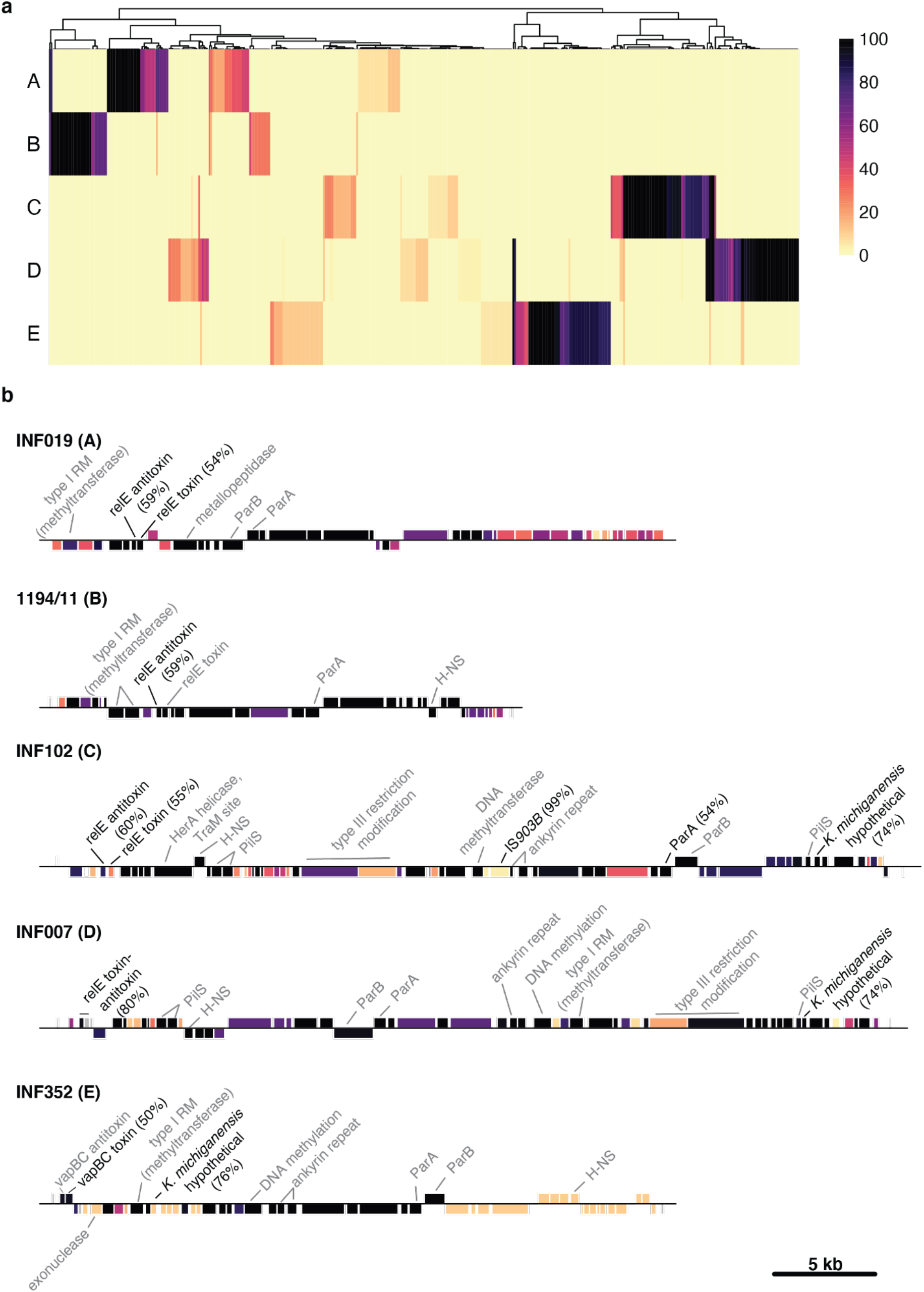
Conservation and function of genes in each phylogroup. **a,** Heatmap showing proportion of plasmids with each gene by phylogroup. Columns are genes, clustered using *hclust;* rows are phylogroups (unclustered). Colour within each cell indicates the proportion of plasmids carrying each gene, as per legend. **b,**Gene maps of one representative plasmid per phylogroup. Genes are indicated by blocks (above line - forward orientation, below line - reverse orientation) and coloured by conservation in their phylogroup. Genes with ≥50% homology to known genes in Enterobacteriaceae are indicated by black lines and text, with gene homology shown in brackets. Genes with detected PFAM domains are indicated by grey lines and text. Details of each gene can be found in **Table S4**.

The vast majority of genes annotated in each linear plasmid were hypothetical proteins and had no close homologs in other Enterobacteriaceae genomes (**Figure 4b**). However, there were few reference plasmid genes (n=55, 20%) for which we were able to obtain some form of functional annotation based on sequence homology or protein domain matches (see **Methods, Table S4**). Most of these annotations were for genes encoding proteins likely relevant to basic plasmid maintenance functions. All five phylogroups carried genes with type II toxin-antitoxin domains (see **Table S4**), which are often found on plasmids and can enable plasmid maintenance by performing post segregational killing of daughter cells that do not carry the plasmid [28]. These systems were core in all phylogroups. Phylogroups A, B, C and D each carried a *relBE* family system (65%-83% homology between variants in phylogroups B-D, A carried a distinct variant), whilst phylogroup E carried a *vapBC* system (**Figure 4b**). These toxin-antitoxin clusters generally had at least one gene of the pair encoding a protein with ≥50% homology to toxin-antitoxin systems found in Enterobacteriaceae (**Figure 4b, Table S4**). Pairs of adjacent genes encoding novel proteins with Pfam matches to the partitioning proteins ParA (PF13614 or PF01656) and ParB (PF18821) were detected as core in each phylogroup (**Figure 4, Table S4**). These likely contribute to control of plasmid segregation into daughter cells [29], homologous sequences were not detected in other Enterobacteriaceae. Sequences with homology to the transcriptional repressor *hns* were identified in all phylogroups except A (**Figure 4b**), however the encoded proteins were highly divergent from one another (27-66%) and the genes were classed as separate gene groups by panaroo (**Table S2**). *Hns* are commonly plasmid-encoded and regulate expression of both plasmid and chromosomally-encoded genes. Proteins with hits to known restriction/modification domains were also identified in all reference plasmids, these are frequently encoded by mobile elements and can function as toxin/antitoxin systems to force maintenance of those elements. Phylogroup A was the only phylogroup in which flagellin genes were identified, in n=7/16 plasmid sequences. One of these was plasmid pBSSB1, and the other six were all linear plasmids from *Salmonella enterica* serovar Senftenberg isolated from Switzerland [10]. Phylogroups C and D both carried three core genes apiece harbouring PilS (type IV pilin) domains (**Figure 4b**), which could potentially function as adhesins.

All five phylogroups differed substantially from one another in their basic characteristics, including plasmid length, TIR length and GC content. Phylogroups D and C had the longest plasmids (medians 40.9 kbp and 42.9 kbp respectively), and phylogroup B the smallest (median 23.7 kbp, **Figure 5a**). We calculated the size of TIRs by aligning the beginning of each plasmid to the reverse complement of itself (see **Methods**). We were able to detect a TIR in 57 of the linear plasmid sequences. Those without a TIR were all identified in publicly available assemblies that were assembled using a variety of methods, and we hypothesise that the lack of TIR sequence is most likely due to incomplete or fragmented assembly of the plasmid, rather than lack of TIR in the sequenced molecules. For plasmids where we performed the assembly in-house, we found that the length of the TIR differed substantially between phylogroups, with phylogroups A and D having the longest TIRs (medians 1168 bp and 1074 bp respectively), whilst phylogroups B, C and E had TIRs of approximately half that length (medians 542 bp, 530 bp and 670 bp respectively, **Figure 5b**). There was a high level of sequence conservation for TIRs within phylogroups, with a median of 89-97% similarity in this region in phylogroups A - D (**Data S1, Figure S2**). Phylogroup E was more diverse with an overall similarity of 65%; however, inspection of the alignments revealed that this phylogroup carried two distinct TIR sequences, with a median of 89-99% TIR sequence identity within each TIR grouping (**Data S1, Figure S2**). Finally, %GC for the linear plasmids was very low in comparison to the normal chromosomal %GC range for Enterobacteriaceae, which is typically ~50% (median 57% for the *Klebsiella* carrying linear plasmids). All linear plasmid phylogroups had %GC <40%, with phylogroup B having the lowest out of all the phylogroups (median 28%, compared to 34-35% for other phylogroups, p<2.5×10^−4^ for all comparisons, Wilcoxon test, **Figure 5c**).

**Figure 5:**
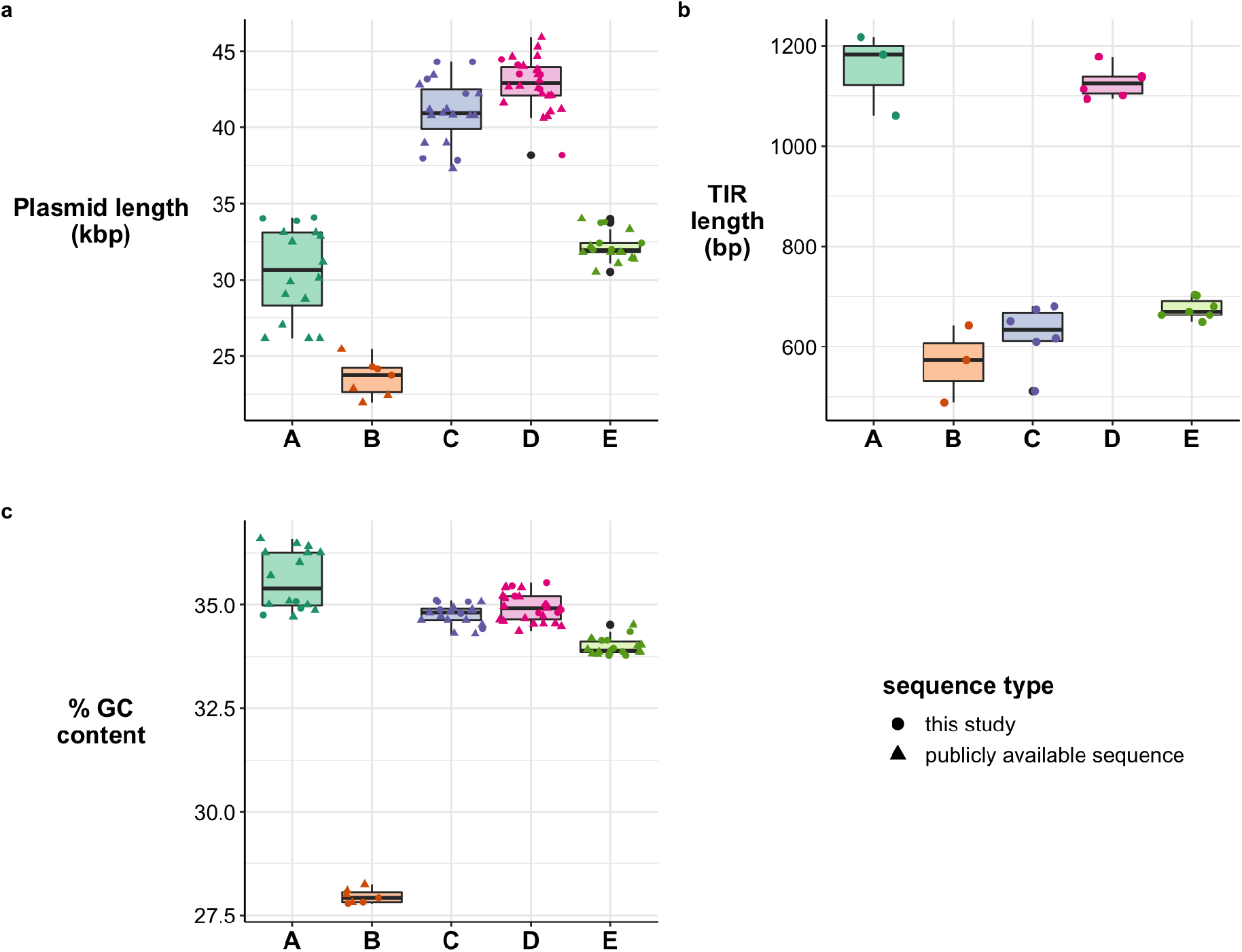
Characteristics of linear plasmid phylogroups. a, Distribution of plasmid lengths in kbp. Boxplots show median (black line), 1^st^ and 3^rd^ quartiles (edges of box), and solid lines give 1.5x the interquartile range. Outliers are shown as black dots. Individual values are shown as coloured dots or triangles, with shape indicating the origin of the sequence as per legend. **b, Distribution of TIR lengths in bp,**as per **(a).**Publicly available sequences are not represented in this plot due to assembly errors in the TIR region. **c, Distribution of GC content,**as per **(a)**.

### *Potential donors of linear plasmids and their stability in* Klebsiella

Given that linear plasmids are rare in *Klebsiella* and have a significantly lower %GC than their host chromosomes, we assume that Enterobacteriaceae are unlikely to be the typical hosts for these plasmids. We used trinucleotide frequencies as a genomic signature to attempt to identify potential original hosts of these plasmids by calculating the distance between our linear plasmids, their *Klebsiella* host chromosomes, and one representative per bacterial species defined in the GTDB (see **Methods**). The 12 *Klebsiella* linear plasmids clustered separately from their corresponding host chromosomes, with a mean distance of 2.3 between the chromosomes and linear plasmids (**Fig S3**). The *Klebsiella* chromosomes were much more similar to each other than the linear plasmids (mean pairwise distance of 0.07 between chromosomal sequences vs 1.27 between pairs of linear plasmid sequences), and clustered closely with other representatives of *Klebsiella* in the GTDB database (nearest neighbour accession GCF_000742135.1, distance 0.09). The linear plasmid from 1194/11 clustered most closely to the Firmicute DUOC01 sp012839065 (accession GCA_012839065.1, distance 1.3). This organism belongs to a strain from the class Thermosediminibacteria, which was detected in a metagenomic sample obtained from an anaerobic digester [30]. The other 11 linear plasmids were their own nearest neighbours (median pairwise distance 1.09), the closest GTDB profile was Proteobacteria isolate Neptuniibacter sp002435145 (accession GCA_002435145.1, mean distance 1.15 to the 11 plasmids). This organism belongs to the order Pseudomonadales, and was detected in a marine environment [31].

To understand whether linear plasmids could be stably maintained within *Klebsiella,* we undertook passage experiments on the 11 *Klebsiella* genomes carrying linear plasmids in our collection. We performed long-read sequencing on all parental isolates (D1), passaged each isolate 10 times (one passage per 24 hour period), and then performed long-read sequencing on the final D10 isolates (see **Methods**). We found that all plasmids, both linear and circular, were maintained in all genomes across 10 passages (**Figure 6**). Linear plasmid copy number was generally estimated at ~1 per cell at both D1 and D10, with the exception of 1194/11 (the only representative of phylogroup B), which had a copy number of 2-4, and two of the phylogroup E plasmids (strains INF345, INF352) with copy number ~2 (see **Figure 6**).

**Figure 6:**
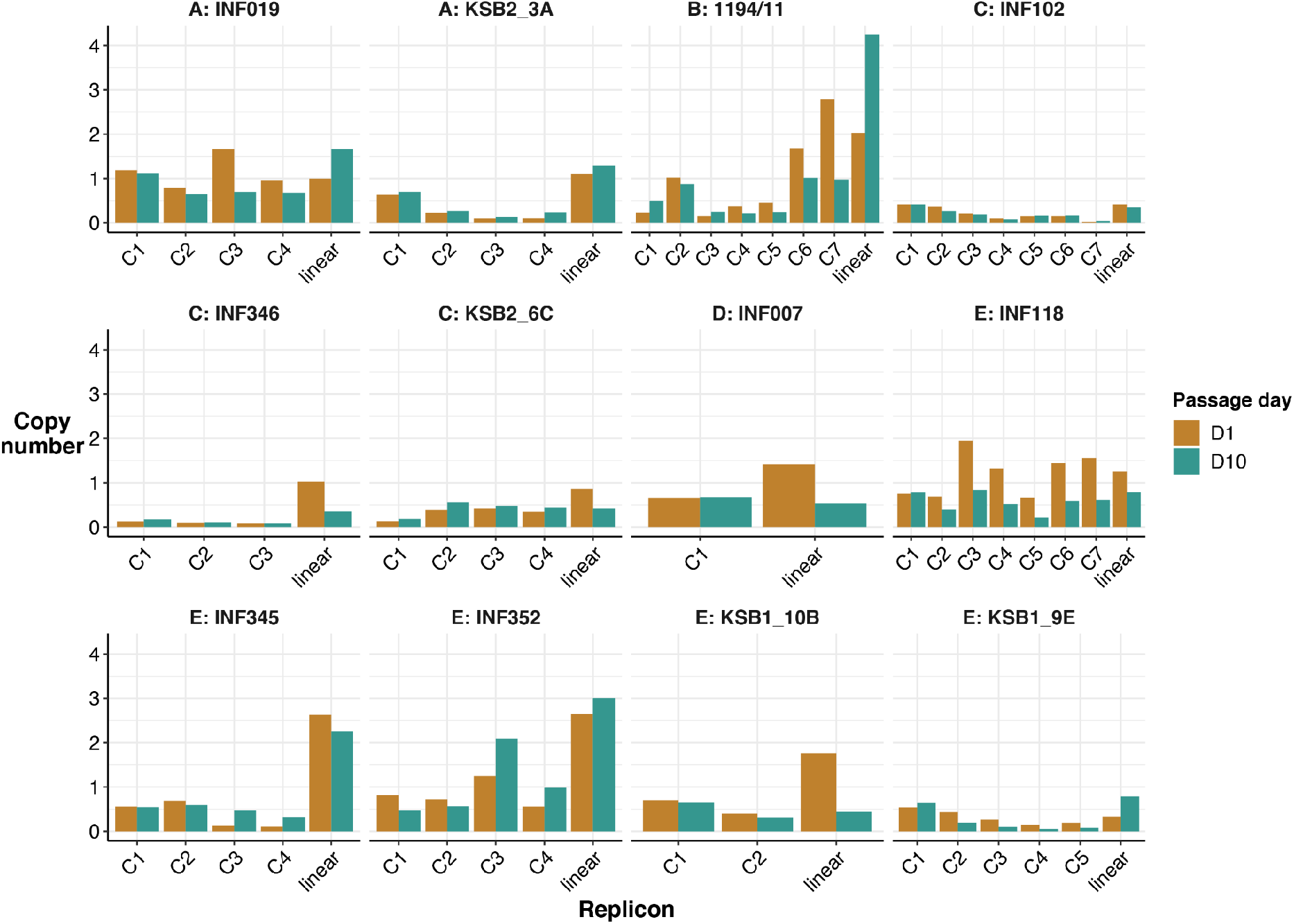
Estimated copy number of all plasmid replicons in each genome. Height of each bar indicates copy number, coloured by passage day as per legend. Each pair of bars represents a plasmid, C[N] indicates a circular plasmid, Linear indicates the linear plasmid in that genome.

## Conclusions

Here we provide the first (to our knowledge) collection of linear plasmids in the *K. pneumoniae* species complex alongside a detailed description of their characteristics. Our data show these plasmids are uncommon in *Klebsiella* and other *Enterobacteriaceae* species, but can be stably maintained and can transfer between distinct *K. pneumoniae* strains (including representatives of the globally-distributed multidrug resistant clones) and other diverse Enterobacteriaceae [4]. The novel *Klebsiella* linear plasmids described here do not carry any known antimicrobial resistance, virulence or metabolic genes; however carriage of a linear plasmid has previously been shown to provide a metabolic advantage for vancomycin-resistant *Enterococcus faecium* in the human gut [3] and to enable flagellar antigen switching in *Salmonella* Typhi. By making freely available these linear plasmid sequences and representative isolates that carry them, we hope to facilitate future research into the function and potential evolutionary or clinical significance of these enigmatic replicons.

## Authors and contributions

Conceptualization, R.R.W, K.L.W and K.E.H; Formal analysis, J.H, H.C, A.T, R.R.W, L.M.J, K.E.H; Methodology, R.R.W, A.T, L.M.J, K.L.W and K.E.H; Software, A.T and R.R.W; Resources, L.C and D.O.G; Visualization, J.H, H.C, R.R.W and K.E.H; Writing - Original Draft, J.H, H.C and K.E.H; Writing - Review & Editing, all authors; Funding acquisition, K.E.H; Project administration, K.E.H.

## Conflicts of interest

The authors declare that there are no conflicts of interest.

## Funding information

K.E.H. was supported by a Senior Medical Research Fellowship from the Viertel Foundation of Australia. This work was supported in part by the Bill & Melinda Gates Foundation OPP1175797. Under the grant conditions of the Foundation, a Creative Commons Attribution 4.0 Generic License has already been assigned to the Author Accepted Manuscript version that might arise from this submission.

---

**Figure S1: Long read alignments to linear plasmids and one representative circular plasmid per *Klebsiella* genome.** The total number of alignments shown is capped at 100 to improve visualisation. The plasmid sequence is the thick black line at the bottom, and reads aligning to the plasmid are shown in green if the alignment starts at the beginning or end of the plasmid sequence, or pink if the alignment starts elsewhere. Segments coloured dotted grey indicate regions of the read that do not align.

**Figure S2: TIR sequence alignments within each phylogroup.** Linear plasmid sequences are clustered by gene content, with the phylogroup indicated by tip colour and coloured as per legend. TIR sequences are aligned within each phylogroup, where each colour represents a different nucleotide as per legend. Colour intensity indicates level of conservation at that position (pale=low; intense=high).

**Figure S3: Cluster dendrogram of trinucleotide frequencies for the 12 linear plasmids and their host *Klebsiella* chromosomes.** Trinucleotide frequencies were clustered using *hclust.* Tips are coloured by phylogroup or chromosome (as per legend).

**Table S1: Details of all linear plasmids described in this study.**

**Table S2: Details of panaroo pangenome analysis.**

**Table S3: Presence/absence of all genes in each linear plasmids.**

**Table S4: Gene annotation details for annotations in representative plasmids, including PFAM hits and hits to *Enterobacteriaceae.***

## Notes

### Competing Interest Statement

The authors have declared no competing interest.

