## Supplementary figures and images for "Linear plasmids in *Klebsiella* and other Enterobacteriaceae"

### Figure S1

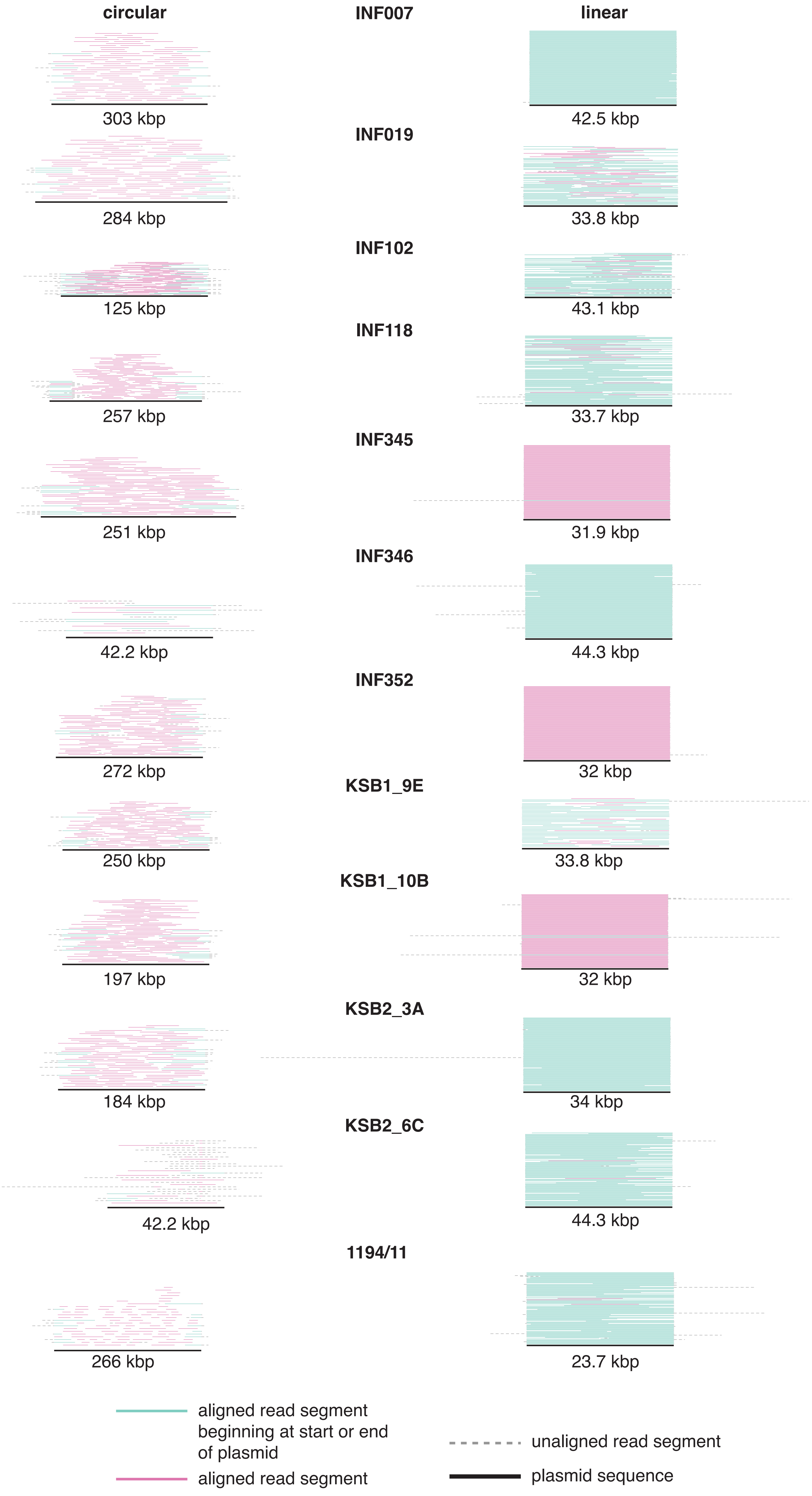

### Figure S2

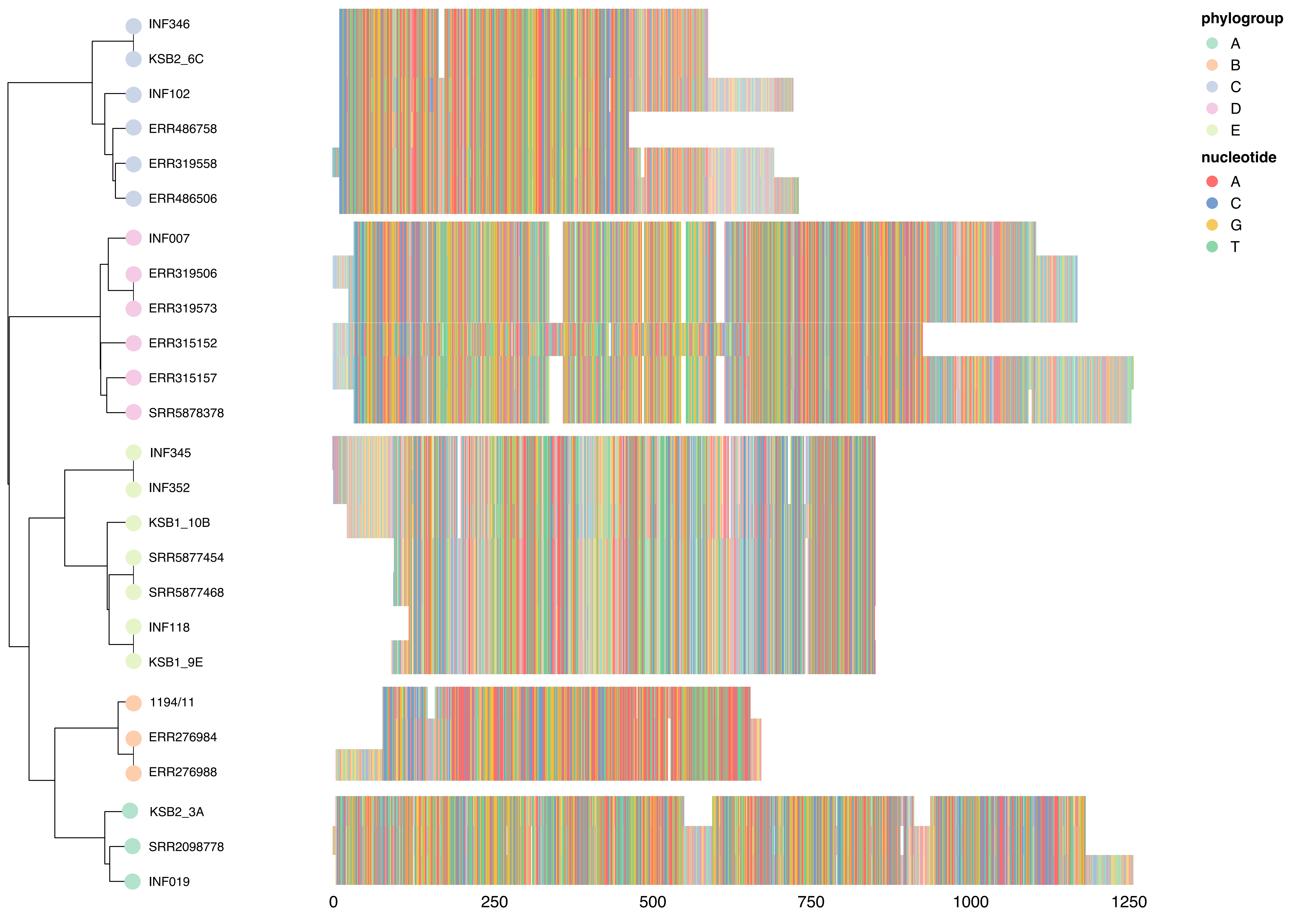

### Figure S3

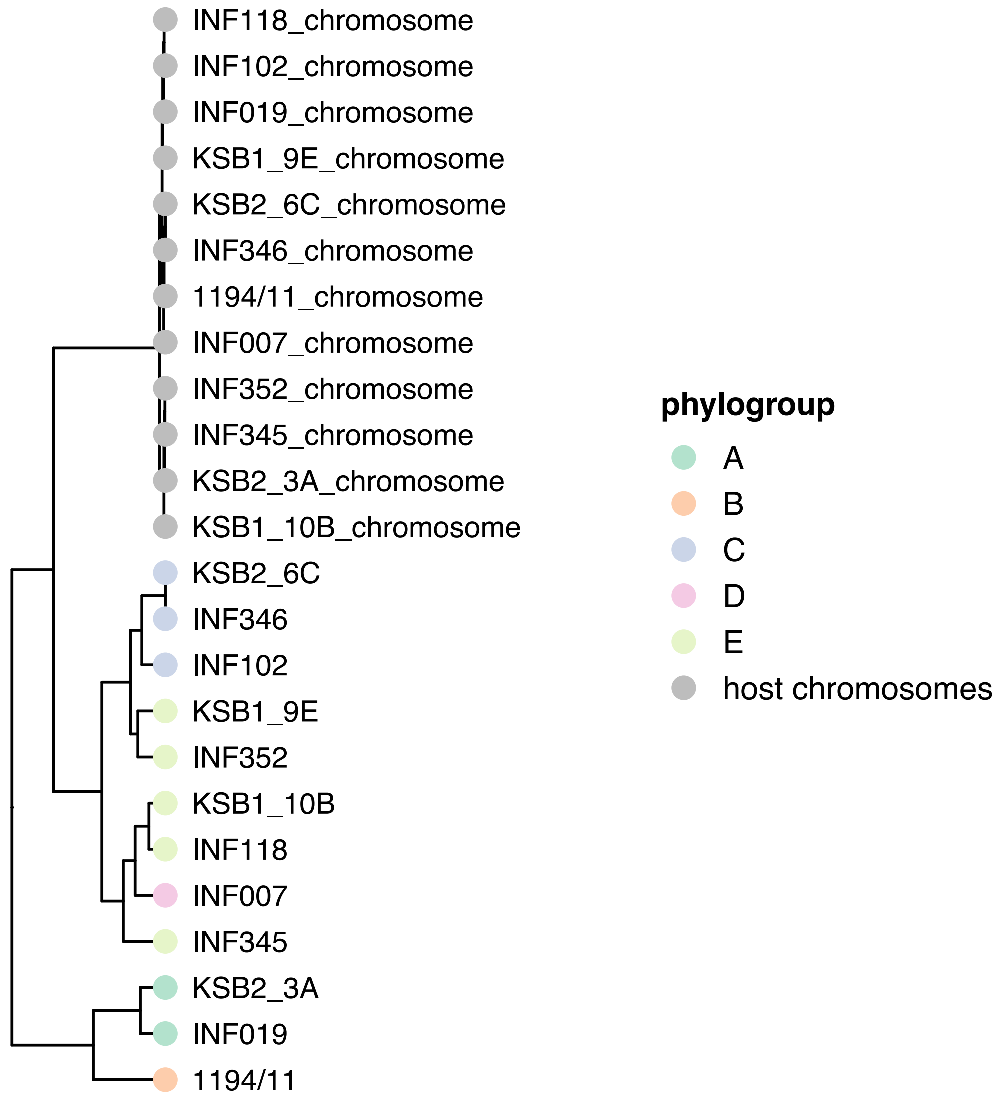
